# Hardy-Weinberg Equilibrium in the Large Scale Genomic Sequencing Era

**DOI:** 10.1101/859462

**Authors:** Nikita Abramovs, Andrew Brass, May Tassabehji

## Abstract

Hardy-Weinberg Equilibrium (HWE) is used to estimate the number of homozygous and heterozygous variant carriers based on its allele frequency in populations that are not evolving. Previously, deviation from HWE in large population databases were investigated to detect genotyping errors, which can result in extreme heterozygote excess (HetExc). However, HetExc might also be a sign of natural selection since recessive disease causing variants are expected to occur less frequently in a homozygous state in the general population, but might reach high allele frequency, especially if they are advantageous, in a heterozygote state. We developed a filtering strategy to detect these variants and applied it on genome data from 137,842 individuals. We found that the main limitations of this approach were quality of genotype calls and insufficient population sizes, whereas population structure and high level of inbreeding could reduce sensitivity, but not precision, in certain populations. Nevertheless, we identified 365 HetExc variants in 326 genes, most of which were specific to African/African American populations (~84.7%). Although the majority of them were not associated with known diseases, or were classified as “benign”, they were enriched in genes associated with autosomal recessive diseases. The resulting dataset also contained two known recessive disease causing variants with evidence of heterozygote advantage in the genes ***HBB*** and ***CFTR***. Finally, we provide *in silico* evidence of a novel heterozygote advantageous variant in the ***CHD6*** gene (involved in influenza virus replication). We anticipate that our approach will allow the detection of rare recessive disease causing variants in the future.

## Introduction

The Hardy-Weinberg Equilibrium (HWE) is an important fundamental principal of population genetics, which states that “genotype frequencies in a population remain constant between generations in the absence of disturbance by outside factors” (Edwards, 2008). According to HWE, for a locus with 2 alleles *A* and *a* with corresponding frequencies p and q, 3 genotypes are possible *AA*, *Aa* and *aa* with expected frequencies p^2^, 2pq, q^2^, respectively (Graffelman et al., 2017). However, various factors, including natural selection, nonrandom mating, genetic drift and gene flow can cause deviations from HWE (Graffelman et al., 2017). Nonrandom mating due to geographical location might be a common cause of deviations from HWE due to heterozygous deficiency in large populations of different ethnicities (Garnier‐Géré and Chikhi, 2013). If a population consists of several subpopulations and individuals randomly mate within, but not between subpopulations, then homozygous alleles in the overall population will be observed more frequently than expected by HWE (the “Wahlund effect”) (Sinnock, 1975). A technical cause of deviations from HWE, sometimes observed in population studies, is sequencing errors (Chen et al., 2017; Graffelman et al., 2017). Previous studies found that variants deviated from HWE mainly due to heterozygote excess (60-69% of the cases) (Chen et al., 2017; Graffelman et al., 2017) and that deviations were 11 times more frequently observed in unstable genomic regions such as segmental duplications and simple tandem repeats (Graffelman et al., 2017), regions that are notoriously prone to sequencing errors. These issues were addressed in the Genome Aggregation Database (gnomAD; release v2.1.1) (Karczewski et al., 2019), the largest publicly available population database at the time of this study (137,842 predominantly healthy individuals from 7 major ethnic populations), which is used in genomic medicine to help prioritise disease causing variants. Variants with extreme heterozygote excess were excluded from gnomAD, whereas those located in repeat regions were marked as dubious.

A factor of deviations from HWE which, to the best of our knowledge, has not yet been investigated on a large scale, is natural selection. Although individuals with known severe pediatric diseases were excluded from gnomAD (Karczewski et al., 2019), some disease causing variants persisted (Tarailo-Graovac et al., 2017). For example, the African population specific (~91% of the carriers) c.20A>T (rs334) variant in the *HBB* gene is a known recessive pathogenic variant which causes sickle-cell disease (MIM:603903) (Ashley-Koch et al., 2000), but it is present in 4 African individuals in a homozygous state (who could be affected by the disease) in gnomAD. Moreover, this variant is present in a heterozygous state in ~9% (1,113/12,482) of African individuals (i.e. unaffected carriers), which is significantly more (~2.5 times) than the expected number (~439 individuals) according to HWE (p-value = 1.38E-07), for that number of homozygous individuals. The presence of a recessive disease causing variant at a high frequency in populations might also be a sign of overdominant selection, i.e. a variant in the heterozygous state provides some advantage to its carriers (Withrock et al., 2015), as is the case for *HBB* c.20A>T which provides carriers protection from malaria (MIM:611162) (Allison, 1954). This example illustrates that variants deviating from HWE due to heterozygote excess might also be recessive disease causing and possibly heterozygote advantageous. Here we developed a variant filtering strategy to detect potential disease causing variants that might deviate from HWE due to natural selection, and applied it on data from the gnomAD database.

## Materials and Methods

### Collecting gene and variant datasets

Initial gene dataset, with disease phenotype and inheritance pattern data from Online Mendelian Inheritance in Man (OMIM) database (Hamosh et al., 2005), was obtained from Gene Discovery Informatics Toolkit (GDIT) (Dawes et al., 2019) and consisted of 19,196 protein coding genes. Population variant data with clinical significance annotation (ClinVar (Landrum et al., 2016)) was obtained programmatically from gnomAD (Karczewski et al., 2019) via API (https://gnomad.broadinstitute.org/api). The database consisted of 137,842 individuals from 7 populations: Non-Finnish European (NFE, n = 64,603), Latino/Admixed American (AMR, n = 17,720), South Asian (SAS, n = 15,308), Finnish (FIN, n = 12,562), African/African American (AFR, n = 12,487), East Asian (EAS, n = 9,977) and Ashkenazi Jewish (ASJ, n = 5,185) (Karczewski et al., 2019). The initial variant dataset consisted of more than 17 million unique variants in 18,214 genes whose names in the GDIT dataset were found in gnomAD.

### Filtering initial variant dataset

Variants which satisfied the following criteria were selected for initial analysis of deviations from Hardy-Weinberg Equilibrium (HWE): **(i)** Variant has to be located in the canonical transcript of a gene (as defined in gnomAD who used GENCODE (Frankish et al., 2019) v19 annotation); **(ii)** Variant has to be located on an autosomal chromosome; **(iii)** Variant has to be protein coding, i.e. has one of the following Variant Effect Predictor (VEP) (McLaren et al., 2016) version 85 consequences: *“transcript_ablation”*, *“splice_acceptor_variant”*, *“splice_donor_variant”*, *“stop_gained”*, *“frameshift_variant”*, *“stop_lost”*, *“start_lost”*, *“transcript_amplification”*, *“inframe_insertion”*, *“inframe_deletion”*, *“missense_variant”*, *“protein_altering_variant”*, *“splice_region_variant”*, *“incomplete_terminal_codon_variant”*, *“start_retained_variant”*, *“stop_retained_variant”*, *“synonymous_variant”*; **(iv)** Variant Allele Frequency (AF) has to be >0.001 in at least one population; **(v)** Variant site has to be covered in at least 80% of the individuals in each of the 7 populations; **(vi)** Variant has to be “PASS” quality in exomes and genomes datasets (if present in both); **(vii)** Variant site must not contain frequent alternative variants that could compromise statistical results of the biallelic HWE test (sum of AFs of all alternative variants seen at the same chromosomal position in the same population has to be <0.001).

### Statistics and measuring deviations from Hardy-Weinberg Equilibrium (HWE)

The original code to measure statistical significance of variant deviation from HWE, developed by Wigginton et al (Wigginton et al., 2005), calculated p-values as the probability of observed sample plus the sum of all probabilities of more extreme cases. However, Graffelman and Moreno (Graffelman and Moreno, 2013) later showed that mid p-value, which is calculated by adding only half of probability of observed sample, had better error rate and statistical power for testing deviations from HWE. Therefore, we used the code created by Wigginton et al, but modified it to return mid p-values. Variants deviating from HWE with mid-p value less than 0.05 were considered to be statically significant. Two sided Fisher’s exact test was used for all other statistical tests unless stated otherwise. Code used in this study can be found at https://github.com/niab/hwe.

### Selecting candidate disease/heterozygote advantageous variants

Variants that satisfied the following criteria were selected into a final set of candidate disease/heterozygote advantageous variant dataset: **(i)** Variant AF has to be <0.1 in each of the 7 populations; **(ii)** Variant must have statistically significant (p-value ≤ 0.05) excess of heterozygotes in at least one population; **(iii)** Variant must have excess of heterozygotes in each population (not required to be statically significant). This filter was added as variants with heterozygote excess in one population but not in the others, might be a result of gene flow; **(iv)** Variant must not be located in a segmental duplication (Bailey et al., 2002) or tandem repeat region (Benson, 1999), which were obtained via UCSC Genome Browser (Haeussler et al., 2019) same as in the Graffelman et al study (Graffelman et al., 2017); **(v)** 50% of variant carriers in the overall population must have Allele Balance (AB) between 0.4 and 0.55 (AB thresholds are justified in the results section). After applying these filters the resulting dataset consisted of 365 unique variants (369 if counted in each population separately) located in 326 genes (Supplementary Table 1).

## Results

After applying initial filters on variant data from 7 major populations (Figure 1A), the resulting dataset consisted of 382,506 unique variants (803,584 if counted in each population separately, Figure 1B) located in 16,871 genes. Exclusion of rare variants (AF < 0.001) from the analysis reduced the possible impact of population size (Figure 1A) on the number of variants analysed (Figure 1B). For example, the Finnish (FIN) populations was ~5 times smaller than the Non-Finnish European (NFE) population (12,562 and 64,603 individuals respectively), but had a similar number of unique variants (85,553 and 92,458 variants respectively). However, population size had a significant effect on the ability of the HWE test to detect Heterozygote Excess (HetExc) deviation in rare variants: the larger the population, the smaller the AF threshold after which statically significant HetExc can be reported. The minimal HetExc AF thresholds (i.e. assuming complete absence of homozygotes) are shown on Figure 1C, note the negative correlation with population sizes shown in Figure 1A.

**Figure 1.**
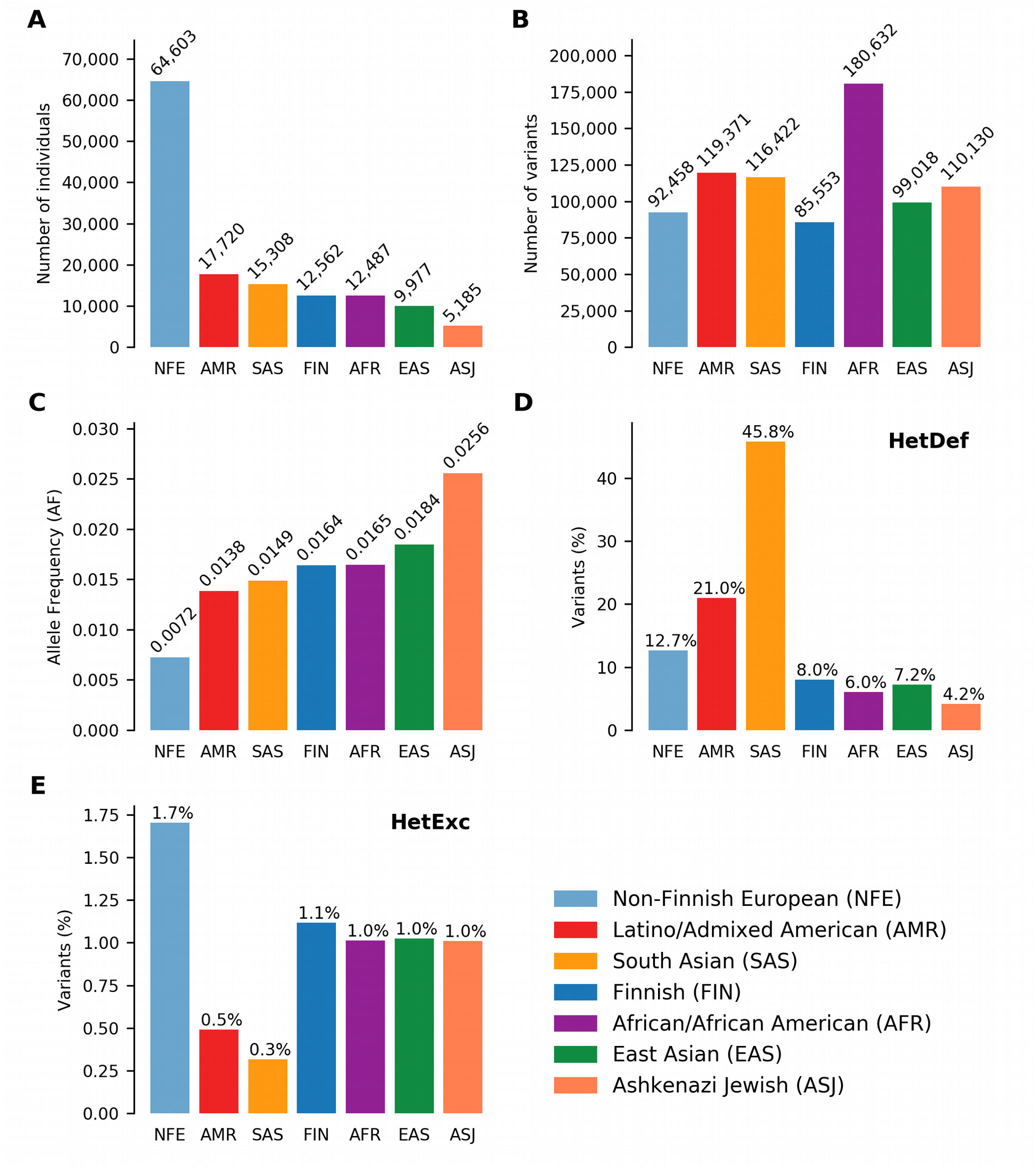
Deviations from Hardy-Weinberg Equilibrium (HWE) in 7 major gnomAD populations. Number of **(A)** individuals and **(B)** variants in each population. **(C)**Minimum variant Allele Frequency (AF) required for statistically significant heterozygote excess according to HWE, in the absence of homozygous individuals in each population. Percentage of variants (raw numbers are shown in **B)** deviating from HWE due to **(D)** heterozygote deficiency or **(E)** heterozygote excess in each population.

Another factor that could affect detection of HetExc variants, was the degree to which HWE assumptions were satisfied in each population. For example the “random mating” assumption would be violated in populations with a high degree of consanguineous marriages or consisting of individuals from several countries, and would result in a higher proportion of variants deviating from HWE due to Heterozygote Deficiency (HetDef) (i.e. “Wahlund effect”). To some degree, all populations deviated more frequently from HWE due to HetDef than HetExc (see Figure 1D,E). The largest proportion of HetDef variants were observed in South Asian (SAS) and Latino/Admixed American (AMR) populations, 45.8% and 21.0% respectively. Consequently, these populations also had the lowest proportion of HetExc variants, 0.3% and 0.5% respectively. The lowest proportion of HetDef variants was observed in the Ashkenazi Jewish (ASJ) population. However, even in this population, variants deviated from HWE due to HetDef ~4 times more frequently than due to HetExc, 4.2% and 1.0% respectively. Interestingly, the African/African American (AFR) population had the second lowest percentage of HetDef variants (6.0%), which outscored the FIN population (8.0%), considered as a homogeneous isolate. The largest proportion of HetExc variants was in the NFE population (1.7%, 1574 variants), which had the smallest AF threshold for HetExc detection (AF = 0.0072, Figure 1C). Despite this, the AFR population still had the largest absolute number of HetExc variants (1,829). Therefore, overall population variant shift from HWE towards HetDef (i.e. the majority of the variants have higher than expected homozygous AF) decreased the number of statistically significant HetExc variants (especially in SAS and AMR), which can also be seen in Figure 2 for relatively rare variants (AF < 0.1).

**Figure 2.**
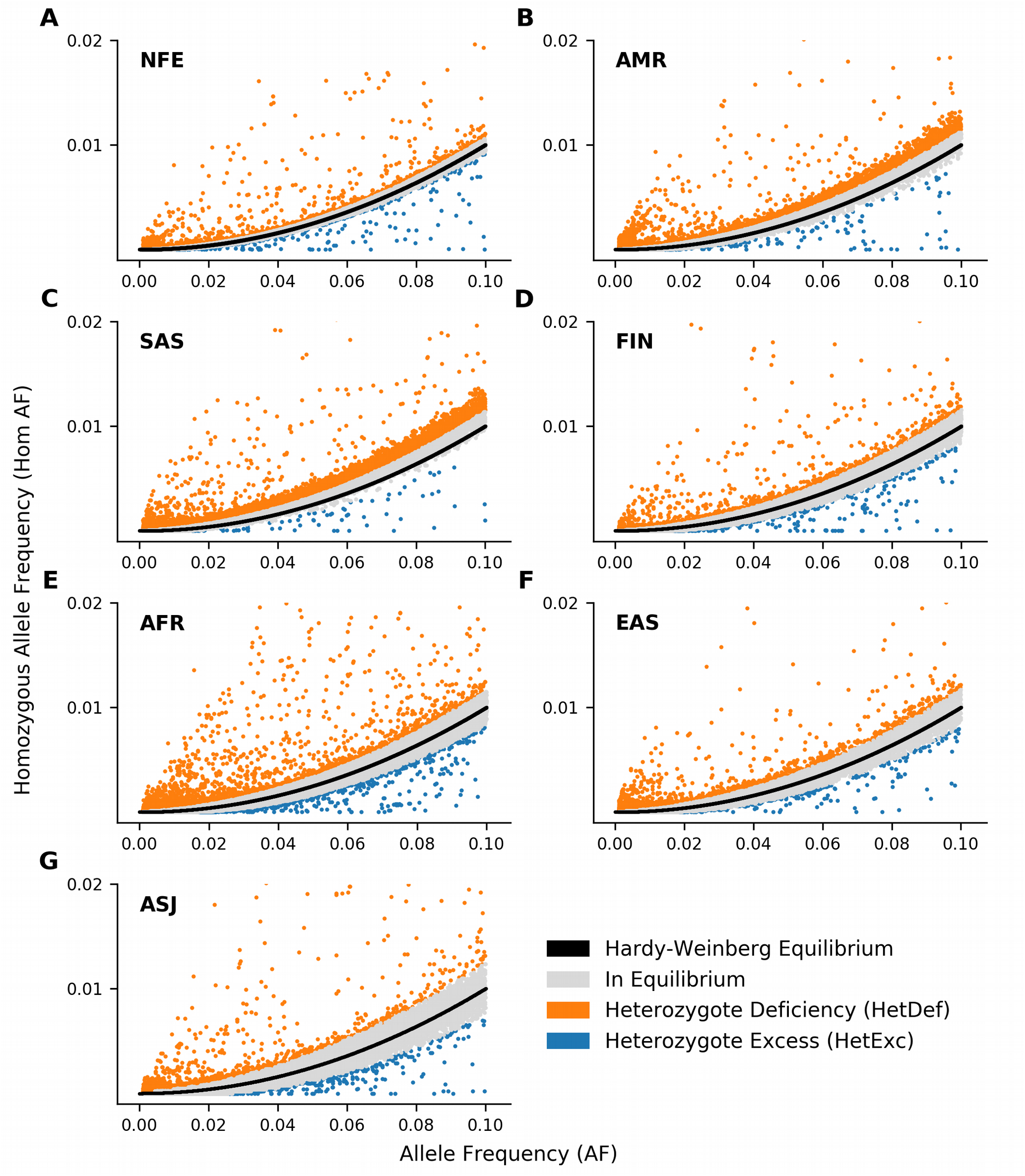
Comparison of observed ratio between variant Alllele Frequency (AF) and homozygous AF with expected ratio according to Hardy-Weinberg Equilibrium (HWE) in 7 major gnomAD populations: Non-Finnish European (**A**, NFE), Latino/Admixed American (**B**, AMR), South Asian (**C**, SAS), Finnish (**D**, FIN), African/African American (**E**, AFR), East Asian (**F**, EAS) and Ashkenazi Jewish (g, ASJ). Black line represents expected ratio between AF and expected homozygous AF according to HWE (i.e. AF^2^). Variants where deviation from HWE are not significant (p-value > 0.05) are shown in gray, whereas those that deviate from HWE due to heterozygote deficiency or excess are shown in orange and blue respectively. Only variants with 0.001 ≤ AF ≤ 0.1 and homozygous AF ≤ 0.02 are shown.

This initial analysis has been performed on variants that remained following the gnomAD sequence quality filtering process and might therefore be assumed to be real. However, variant databases are known to contain errors which could give a significant HetExc signal. To explore this we have developed a set of more stringent filters. In particular, variant properties that could produce a false positive HetExc signal were investigated. For this analysis, variants present in multiple populations were counted once, and variants with AF > 0.1 in at least one population were excluded. A high AF threshold was used to exclude common variants that are unlikely to be recessive disease causing, but retain relatively common variants that still might be real (e.g. *HBB* c.20A>T (Ashley-Koch et al., 2000), AFR AF = ~0.045). At this stage, only variants which had an excess of heterozygotes in all populations, and were statistical significant in at least one population were classified as HetExc.

Firstly, to investigate the correlation between HetExc and chromosomal regions prone to sequencing errors, variants were divided into 3 groups: (i) “segmental duplication” (3,762), (ii) “tandem repeat” (1,588) and (iii) all others named “Ref” (54,897). HetExc variants were significantly more frequent in both “tandem repeat” (Fold Enrichment (FE) = ~2.1, p-value = ~0.001) and “segmental duplication” (FE = ~3.3, p-value = 2.0E-21) groups than in the “Ref” group (Figure 3A). Therefore, HetExc of variants located in segmental duplications and tandem repeats might be a result of genotyping errors and were excluded from further analysis.

**Figure 3.**
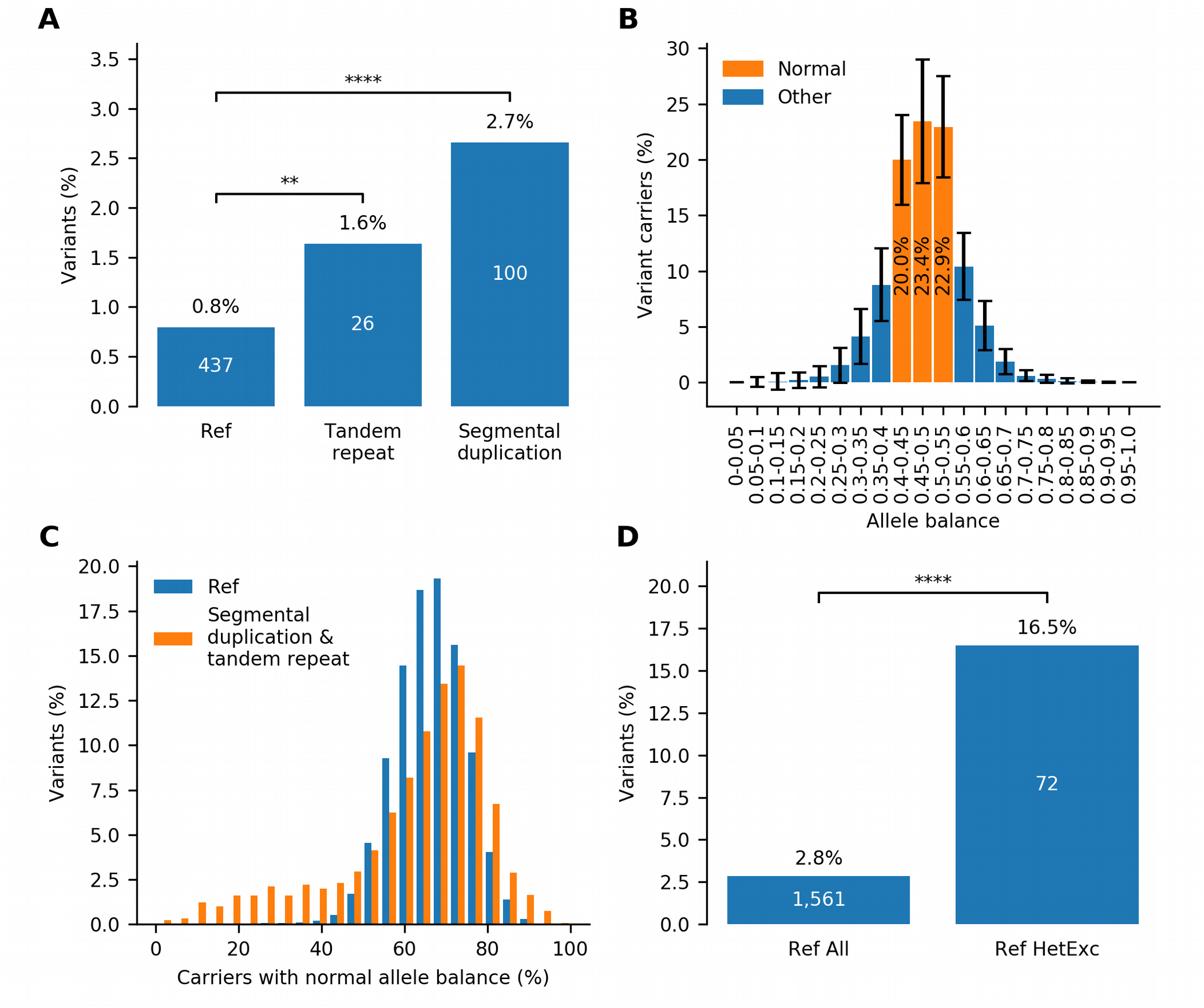
Impact of tandem repeats, segmental duplications and allele balance on the probability of variant deviation from Hardy-Weinberg Equilibrium (HWE) due to heterozygote excess (HetExc). (A) Percentage of variants deviating from HWE due to HetExc which are located in tandem repeat, segmental duplication regions or reference (“Ref”) group (i.e. all other regions). (B) Distribution of Allele Balance (AB) between variant carriers in variants from “Ref” group (error bars indicate standard deviation). For each variant these statistics are aggregated into a single metric that represents cumulative percentage of Variant Carriers with Normal (0.4-0.55) Allele Balance (VCNAB, e.g. 20.0% + 23.4% + 22.9% = 66.3%). (C) Distribution of variants with various VCNAB percentages in “Segmental duplication & tandem repeat region” and “Ref” groups. (D) Percentage of variants with VCNAB < 50% in the whole “Ref” group and a subset of variants with statistically significant excess of heterozygotes in “Ref” group.

Secondly, to investigate the correlation between HetExc and Allele Balance (AB), which is a known indicator of systematic genotyping errors (Muyas et al., 2019), the AB profile of an average gnomAD variant was required. In gnomAD, variant AB data is stored as a number of variant carriers (converted to percentages here) in 20 AB groupings (from 0 to 1, 0.05). Figure 2B shows the distribution of AB between variant carriers in variants from the “Ref” group. For an average variant, the majority of variant carriers (66.3%) had an AB between 0.4 and 0.55 and were named “Normal” here, because it was close to the expected normal 0.5 ratio for heterozygous variants. To aggregate variant AB data into a single numeric metric, it was measured as percentage of Variant Carriers with “Normal” Allele Balance (VCNAB). The combined “segmental duplication” and “tandem repeat” group had more variants with high and low VCNAB (80% Confidence Interval (CI) = 34.0-80.1%) than the “Ref” group (80% CI = 55.6-76.9%), which correlated with previous results showing that variants in repetitive regions were more prone to genotyping errors (Figure 3C). Minimal VCNAB threshold for “PASS” quality variants was defined by a lower bound fraction of 95% CI calculated for variants from the “Ref” group (CI = 49.4-82.1%), and was rounded to 50% (i.e. half of the variant carriers must have AB in the range 0.4-0.55). Only 2.8% of variants in the “Ref” group would not pass this filter, but among HetExc variants, the fail rate would be ~5.8 times higher (p-value = 7.0E-29, Figure 3D). Therefore, variants with low AB (VCNAB < 50%) might be enriched with genotyping errors and were also excluded from further analysis.

Finally, HetExc variants which were not located in segmental duplication or tandem repeat regions and had VCNAB ≥ 50% were selected as candidate recessive disease causing genes (Supplementary Table 1) and were then compared with a group of variants that survived the same filtering process, but did not have an excess of heterozygotes (HetExc-). The HetExc- and HetExc groups consisted of 52,947 and 365 variants (82,066 and 369 if counted in 7 ethnic populations separately, Figure 4A) in 12,714 and 326 genes respectively. Both HetExc- and HetExc groups had similar proportions of missense and synonymous variants (Figure 4B). Most of the HetExc variants were present in African/African American populations (309/365, ~84.7%), which was significantly more than expected (FE = 1.6, p-value = 6.4E-10) based on the proportion in the HetExc-group (27,561/52,947). All other populations had significantly less than expected HetExc variants (East Asian FE = ~0.6 p-value = ~0.008, all other populations FE ≤ 0.25, p-value ≤ 1.0E-10).

**Figure 4.**
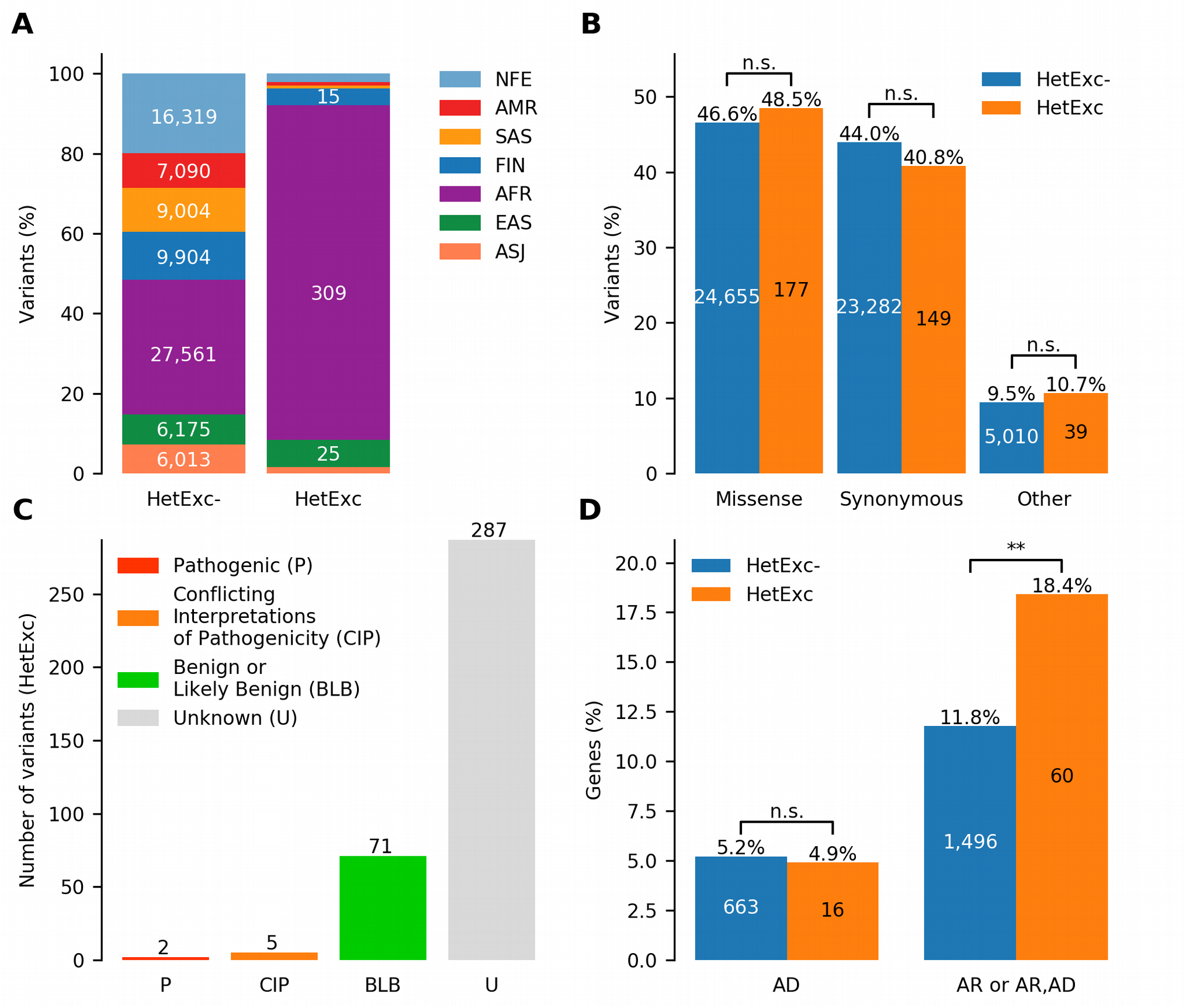
Potential recessive disease causing variants identified by statistically significant deviation from Hardy-Weinberg Equilibrium (HWE) due to excess of heterozygotes (HetExc). **(A)** Distribution of variants deviating and not deviating from HWE due to excess of heterozygotes (HetExc and HetExc-respectively) in 7 major gnomAD populations. **(B)** Proportions of missense, synonymous and other protein coding variants in HetExc and HetExc-datasets. **(C)** ClinVar clinical significance status (e.g. pathogenic/benign) of HetExc variants **(D)** Known disease associated genes with at least one variant in HetExc and HetExc-datasets grouped by inheritance pattern: Autosomal Dominant (AD), Autosomal Recessive (AR) or both.

To understand which of the HetExc candidate recessive disease causing variants were already known, their clinical significance status in the disease variant database (ClinVar (Landrum et al., 2016)) was analysed. The majority of HetExc variants (287/365, 78.6%) were not present in ClinVar, whereas the majority of those that were present in ClinVar (71/78, 91.0%) had “Benign” or “Likely benign” status (Figure 4C). The only two variants with “Pathogenic” status were c.20A>T (rs334) in *HBB* (causes recessive sickle cell disease MIM:603903; carriers are protection from malaria, MIM:611162) and c.1521_1523delCTT (rs1801178) in *CFTR* (causes recessive cystic fibrosis disease, MIM:219700; hypothesised to be protective from cholera (Rodman and Zamudio, 1991) or tuberculosis (Bosch et al., 2017)). However, genes with at least one HetExc variant were significantly more frequently associated with known Autosomal Recessive (AR) diseases than genes containing only HetExc- variants (FE = ~1.6, p-value = ~0.0025, Figure 4D). HetExc variant enrichment in known AR genes adds evidence that some of the selected variants might deviate from HWE due to natural selection and have some association with disease.

A new version of the gnomAD database was released (v3, containing only whole genomes not exomes; 71,702 samples mapped to build GRCh38) during the preparation of data for this study, which contained a larger AFR population (~1.7 times larger; 21,042 individuals, all other populations were smaller than in gnomAD v2.1.1). We re-analysed 309 AFR HetExc variants from gnomAD v2.1.1 and found that only 43 (~14%) of them were also HetExc in gnomAD v3 (annotated in Supplementary Table 1, chromosomal coordinates were mapped with LiftOver (Haeussler et al., 2019)), which highlights potential differences in the inclusion criteria of individuals or genotype calling procedures between the two releases of the database.

## Discussion

Analysis of deviations from the Hardy-Weinberg Equilibrium (HWE) on a large genomic dataset has shown that all populations, but especially South Asian (SAS) and Latino/Ad-mixed American (AMR), were more frequently deviating from HWE due to heterozygote deficiency (HetDef) than heterozygote excess (HetExc). A Higher rate of HetDef variants in SAS and AMR populations is in line with previous reports (Gazal et al., 2015; Chen et al., 2017), possibly due to the large number of consanguineous marriages in these regions (e.g. 38% of SAS population in the Exome Aggregation Consortium (ExAC) (Lek et al., 2016)). However, our findings that HetDef is a major cause of deviations from HWE in all populations is contrary to previous studies performed by (Graffelman et al., 2017) and (Chen et al., 2017), who used more strict p-value thresholds (0.001 and 0.0001, respectively) and reported that deviations from HWE were more frequently observed due to HetExc. However, previous studies were focused on error detection in older and smaller datasets, some of which were corrected in gnomAD. Graffelman et al. analysed 104 Japanese individuals in the 1000 Genomes database (The 1000 Genomes Project Consortium, 2015), and in these circumstances the minimal statistically significant HetExc AF threshold (0 homozygous and p-value < 0.001) was ~0.23 (Graffelman et al., 2017). Only 11/382,506 variants analysed in our study were that frequent and had no homozygous individuals reported, 9 of which were located in segmental duplication or tandem repeat regions. A higher rate of HetExc variants in these regions, as well as those that had low allele balance, observed in this study, correlates with previous work (Graffelman et al., 2017; Muyas et al., 2019). Chen et al. analysed “open reading frame” genes and selected only 1 variant per gene where AF was closer to 0.50 (584 variants in total) in ExAC (60,706 individuals) (Chen et al., 2017). However, this approach resulted in exclusion of rare variants that were analysed in this study and might be more affected by the Wahlund effect (i.e. more likely to be HetDef). Moreover, some of the HetExc variants detected in previous studies were marked as non-pass quality or were no longer HetExc in gnomAD, possibly due to differences in variant filtering and genotype calling procedures. For example, c.1801C>T (rs1778112) in ****PDE4DIP**** was present in the heterozygous state in ~91% of individuals, and was never observed as homozygous in the 1000 genomes database, but was non-pass quality in gnomAD. Another example, the ****BRSK2**** variant c.551+6delG (rs61002819) was HetExc in ExAC (p-value = 1.9E-15), but not in gnomAD (p-value = 0.13). Therefore, a higher rate of HetDef variants in our study could be explained by a larger population size and a different variant dataset, as well as improvements in variant filtering and genotype calling procedures.

Analysis of HetExc variants (Supplementary Table 1), selected as recessive disease causing candidates, led to somewhat contradictory results, which should be interpreted with caution. Enrichment of HetExc variants in the African/African American (AFR) population was unexpected, and might indicate more extensive natural selection or be a sign of systematic genotype errors in this population. Enrichment of HetExc variants (70/365) in genes associated with known autosomal recessive diseases supports the hypothesis that some of these variants could be recessive disease causing, whereas the presence of a large proportion of synonymous variants (36/70) and the assigned “Benign” or “Likely benign” status of the majority of the known variants (51/70 in CinVar, 46/51 were “Benign” or “Likely benign”) in this group provides evidence against it. Moreover, despite applying our extensive filtering strategies, many of the HetExc variants might still be deviating from HWE due to genotype errors or by chance due to insufficient population size. The latter might be an explanation for some AFR variants that were HetExc in gnomAD v2.1.1, but not in the new v3 release, which had a larger AFR population. However, the c.1521_1523delCTT (rs1801178) variant in ***CFTR*** was also more frequently observed in the homozygous state in gnomAD v3 (4/32,299 homozygous individuals, AF = ~0.014) than in v2.1.1 (1/64,603 homozygous individuals, AF = ~0.012), although Non-Finnish European (NFE) population was ~2 times smaller. Therefore, the difference between homozygote numbers in gnomAD v2.1.1 and v3 might also be explained by other factors, such as differences between genotype calling procedures used on whole exome and genome data.

Nevertheless, the presence of known pathogenic and heterozygote advantageous variants such as c.20A>T in ***HBB*** and c.1521_1523delCTT in ***CFTR*** suggests that some of the other 363 HetExc variants might also follow the same pattern. Specifically, we would like to highlight the ***CHD6*** gene variant c.7210G>C (rs61292917), which was HetExc in both versions of the gnomAD database and was predicted to be deleterious by *in silico* tools (SIFT = 0 (Ng and Henikoff, 2003); PolyPhen-2 = 0.961 (Adzhubei et al., 2013)). Moreover, it was more frequently (FE = 5.21, p-value = 1.19E-04) seen among African than African American populations in the 1000 genomes database (Supplementary Table 2), similar to the c.20A>T variant in ***HBB*** (FE = 3.42, p-value = 1.49E-05), which suggests that these variants might be under purifying selection in populations which moved out of Africa (i.e. they might be disease causing, but advantageous only in Africa, which is known in the case of ***HBB*** c.20A>T case). Although ***CHD6*** is not yet linked with any disease, it is known to play a role in the influenza virus replication process (Alfonso et al., 2011; Alfonso et al., 2013). Interestingly, c.7210G>C has a much lower AF in the African population (AF = 0.066), than c.20A>T (AF = 0.120) in the 1000 genomes database. Considering ***CHD6*** is extremly intolerant to variation (probability of Loss-of-Function Intolerance (pLI) = 1 (Lek et al., 2016), missense Z-score = 4 (Samocha et al., 2014)), c.7210G>C is more enriched in the African population compared with c.20A>T (i.e. possible due to stronger purifying selection), this suggests that c.7210G>C might be disease causing even in the heterozygous state.

Our study highlighted that the usage of HWE to detect candidate recessive disease causing variants is mainly limited by both the quality of genotype calls and the size of available exome/genome population variant data, whereas absence of information about sub-populations (e.g. Africans and African Americans) and a high level of inbreeding (e.g. SAS) could reduce sensitivity, but not precision, of the approach in certain populations. We anticipate that improvements in sequencing technologies and variant filtering software should reduce the number of false positive HetExc variants in the future. In fact, false positive HetExc variants that survived our strict quality filters, might aid the development of more efficient sequencing filtering strategies by helping to understand new patterns of genotype errors. The size of the largest population analysed in this study (NFE = 64,603 individuals) allowed us to detect statistically significant HetExc only amongst variants with AF ≥ ~0.0072 (~39% of 67,212 variants with AF = 0.001-0.1). Consequently, some common recessive disease causing variants were missed even if homozygous individuals were completely absent in the population. For example, HetExc of the c.448G>C (rs1800546) variant in ***ALDOB*** (causes recessive hereditary fructose intolerance) was not statistical significant (p-value = ~0.3), despite being observed in the heterozygous state in 627 NFE individuals (AF = ~0.005). As the number of sequenced exomes and genomes is rapidly growing, this problem may soon be addressed. Indeed, the United Kingdom National Health Service is planning to sequence 1 million genomes in the next 4 years with a wider ambition to increase this number to 5 million^1^. If the NFE population was 1 million, then the AF threshold would drop to ~0.0018 (~76% of 67,212 variants with AF = 0.001-0.01), whereas with 5 million individuals it would be possible to detect statistically significant HetExc in all variants with AF ≥ ~0.0008. Therefore, it might be possible to use HWE to detect rare recessive disease causing variants in the near future.

## Conclusion

In this study, we explored the use of HWE to identify recessive disease causing candidate variants in a large mainly healthy population database by developing a bespoke filtering strategy to detect variants where an excess of heterozygotes in a population could be a result of natural selection. Overall, this approach showed potential, especially for the AFR population, successfully identifying some variants in recessive diseases that are known to be heterozygote advantageous, and providing novel candidates for further investigation. A natural progression of this work would be validation of genotype calls of HetExc variants to understand possible causes of genotype errors and analysis of the biological effect of true positive HetExc variants to determine their potential health implications. We also anticipate that this approach will become more robust in the future as the size and quality of available genomic data increases.

## Author Contributions

NA, MT, AB conceived and designed the research. NA executed the analysis. NA, MT performed the primary writing. MT, AB supervised all aspects of the research, reviewed and edited the manuscript.

## Funding

This work was supported by the Engineering and Physical Sciences Research Council [EP/N509565/1]. MT was funded by the Newlife Foundation (grant #14-15/15).

## Conflict of Interest

The authors declare that the research was conducted in the absence of any commercial or financial relationships that could be construed as a potential conflict of interest.

## Acknowledgments

The authors would like to acknowledge the support of the Manchester Academic Health Science Centre. The authors would like to thank the Genome Aggregation Database (gnomAD) and the groups that provided exome and genome variant data to this resource. A full list of contributing groups can be found at https://gnomad.broadinstitute.org/about.

## Supplementary Material

Supplemental data includes two tables:

Supplementary Table 1. Dataset of variants deviating from Hardy-Weinberg Equilibrium due to heterozygote excess (HetExc).

Supplementary Table 2. Statistical comparison of Allele Frequencies of heterozygote excess (HetExc) variants in *HBB* and *CHD6* genes between African and African American population in the 1000 Genomes database.

## Footnotes

1 https://www.gov.uk/government/news/matt-hancock-announces-ambition-to-map-5-million-genomes

